# Two epidemics, one genotype, different outcomes: evolutionary changes of Avian Influenza H5N1, genotype EA-2024-DI

**DOI:** 10.64898/2026.05.25.727580

**Authors:** Bianca Zecchin, Isabella Monne, Marta Dianati, Alessio Bortolami, Enrico Savegnago, Erga Shkodra, Sandra Revilla-Fernándezd, Mieke Steensels, Steven Van Borm, Emiliya Ivanova, Ivana Rončević, Vladimir Savić, Alexander Nagy, Charlotte K. Hjulsager, Casper Thorup, Lars E. Larsen, Imbi Nurmoja, Ari Kauppinen, Niina Tammiranta, Francois-Xavier Briand, Beatrice Grasland, Ann Kathrin Ahrens, Anne Pohlmann, Anne Günther, Timm Harder, Peter Malik, Laura Garza Cuartero, Svetlana Cvetkova, Juris Ķibilds, Žanete Šteingolde, Egidijus Pumputis, Simona Pileviciene, Chantal J. Snoeck, Manon Bourg, Oxana Groza, Beatriz Bellido Martin, Ron Fouchier, Sanne Thewessen, Oanh Vuong, Monika Ballmann, Marc Engelsma, Cathrine Arnason Bøe, Anna Pikula, Krzysztof Śmietanka, Margarida Dias Duarte, Margarida Henriques Mourão, Iuliana Onita, Dejan Vidanovic, Zuzana Dirbakova, Martin Tinak, Brigita Slavec, María José Ruano, Maite Barrios, Fereshteh Banihashem, Caroline Bröjer, Stina Hedblom, Siamak Zohari, Claudia Bachofen, Ashley C. Banyard, Holly A. Coombes, Ben Clifton, Benjamin C. Mollett, Joe James, Michael J. McMenamy, Robyn McKenna, Ken Lemon, Calogero Terregino, Alice Fusaro

## Abstract

Since 2020, high pathogenicity avian influenza H5Nx viruses of clade 2.3.4.4b have become enzootic in Europe, causing recurrent epidemic waves characterized by extensive reassortment events. Here, we describe the emergence of a single high-fitness genotype (EA-2024-DI) that has driven two consecutive waves, evolving into distinct sub-lineages. While its circulation is ongoing, during the 2025-2026 wave it caused an unprecedented number of cases in wild birds. Using phylodynamic analyses of a large dataset of genomic sequences, we compared the spatial diffusion and host transmission pattern of the EA-2024-DI sub-lineages across the three most recent epidemic waves (2023-2024, 2024-2025 and 2025-2026). We show that the genotype has persisted over time and has spread primarily through wild Anseriformes, but with a marked change in the transmission patterns between the different waves and a shift in the epicenter from Eastern to Central Europe, the latter having emerged as an important hub for virus diffusion throughout Europe. Our results reveal a recent increase in the frequency of viruses from wild and domestic mammals carrying mutations enhancing virus replication in mammalian hosts, highlighting the importance of proactive monitoring of this group of hosts to better understand its role in the virus ecology and evolution.

## Introduction

Since 2016, Europe has experienced repeated incursions of high pathogenicity avian influenza viruses (HPAIV) of the clade 2.3.4.4b H5Nx subtype^1^. The epidemic waves of 2016-2018 and 2020-2021 were driven by the H5N8 subtype. At the beginning of the 2020-2021 epidemic, a H5N1 reassortant, termed genotype EA-2020-C, emerged in Europe and continued to circulate at low levels throughout the entire epidemic wave. Not only did this genotype become dominant at the beginning of the 2021-2022 epidemic wave in Europe, it also spread from Europe to Africa and Asia, and crossed the North Atlantic, marking the beginning of its dissemination across the Americas, reaching geographic areas that had not previously been affected^2,3^. Since the 2021-2022 epidemic, the virus has become enzootic in European wild bird populations, resulting in year-round circulation, increased environmental persistence and an expanded host range^4^. Its extensive circulation in wild birds has favored reassortment with co-circulating non-notifiable avian influenza viruses, resulting in over 100 distinct H5Nx genotypes in Europe over the past five years. This genomic plasticity has enabled the virus to assemble host-adapted genome constellations, facilitating the colonization of novel avian populations and their respective ecological niches. A striking example is genotype EA-2022-BB, which emerged by acquiring gene segments from H13-subtype viruses enzootic in Charadriiformes^4^. This genetic constellation enabled the virus to efficiently infect and establish dynamic infection cycles within species of this Order.

Conversely, genotypes emerging from reassortment events with non-notifiable AIVs in Anseriformes appear to be commonly detected within species of this host order, and have been frequently linked to outbreaks in domestic birds^5^.

Despite the significant number of H5N1 genotypes that have circulated since H5N1 EA-2020-C, the emergence of the EA-2024-DI genotype appears to have brought about a shift in the evolutionary trajectory of these viruses. Since October 2024, virus evolution appears to have been primarily driven by the gradual accumulation of mutations through genetic drift rather than the dynamic reassortment events observed in previous years. This has led to the diversification of the EA-2024-DI genotype into multiple sub-lineages (EA-2024-DI.1, EA-2024-DI.2, EA-2024-DI.2.1). Detection of this genotype has been dominant across both wild birds and poultry with only occasional novel reassortment events being detected.

Furthermore, since September 2025 mainland Europe has experienced an unprecedented number of outbreaks among wild birds, with numbers reaching six times those recorded during the previous two seasons^6^. Interestingly, significant mortality events have been observed in common cranes^7^ following the Western European Flyway from Scandinavia to the Iberian Peninsula as well as an unprecedented number of detections in wild waterfowl.

To better understand the evolutionary changes and the dominance of sub-lineage EA-2024-DI.2.1, which is responsible for the unprecedented number of cases observed during the current 2025–2026 epidemic, we analysed all available full-length genomic sequences of the EA-2024-DI genotype collected since December 2023. Using both discrete and continuous phylogeographic approaches, we compared the geographic spread and host diffusion patterns of this genotype across the 2023–2024, 2024-2025 and 2025–2026 epidemic waves.

## Results

### Geographic and host distribution of genotype EA-2024-DI and its sub-lineages

Genotype EA-2024-DI consists of five gene segments (PA, HA, NP, NA, and M) derived from the H5N1 EA-2021-AB genotype^4^. The remaining three segments (PB2, PB1, and NS) originated from reassortment events most likely with AIVs circulating in wild Anseriformes in Eurasia.

The EA-2024-DI genotype was first identified in mid-December 2023 in wild Anseriformes in Moldova, Poland, and Romania. During the 2023–2024 epidemic, it co-circulated with other genotypes and was detected in fifteen different host species, mainly (>95%) wild Anseriformes and domestic birds, across ten countries, mostly located in Eastern Europe (Austria, Croatia, the Czech Republic, Germany, Hungary, Italy, Moldova, Poland, Romania, Slovakia). From October 2024 it became the predominant genotype in Europe and diversified into two genotype EA-2024-DI sub-lineages, EA-2024-DI.1 and EA-2024-DI.2 (Figures 1 and 2A).

**Figure 1.**
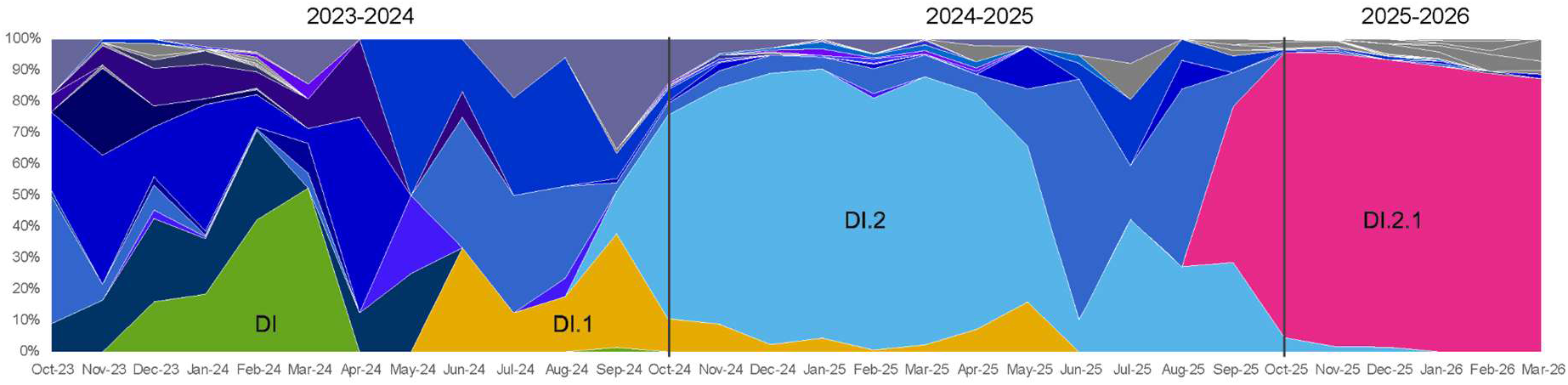
Distribution of genotypes across the three epidemic waves. Genotype EA-2024-DI is highlighted in green, sub-lineages EA-2024-DI.1 in yellow, EA-2024-DI.2 in light blue, and EA-2024-DI.2.1 in pink. Major genotypes recognized by Genin2 are shown in different shades of blue and purple, while minor genotypes are coloured in grey.

**Figure 2.**
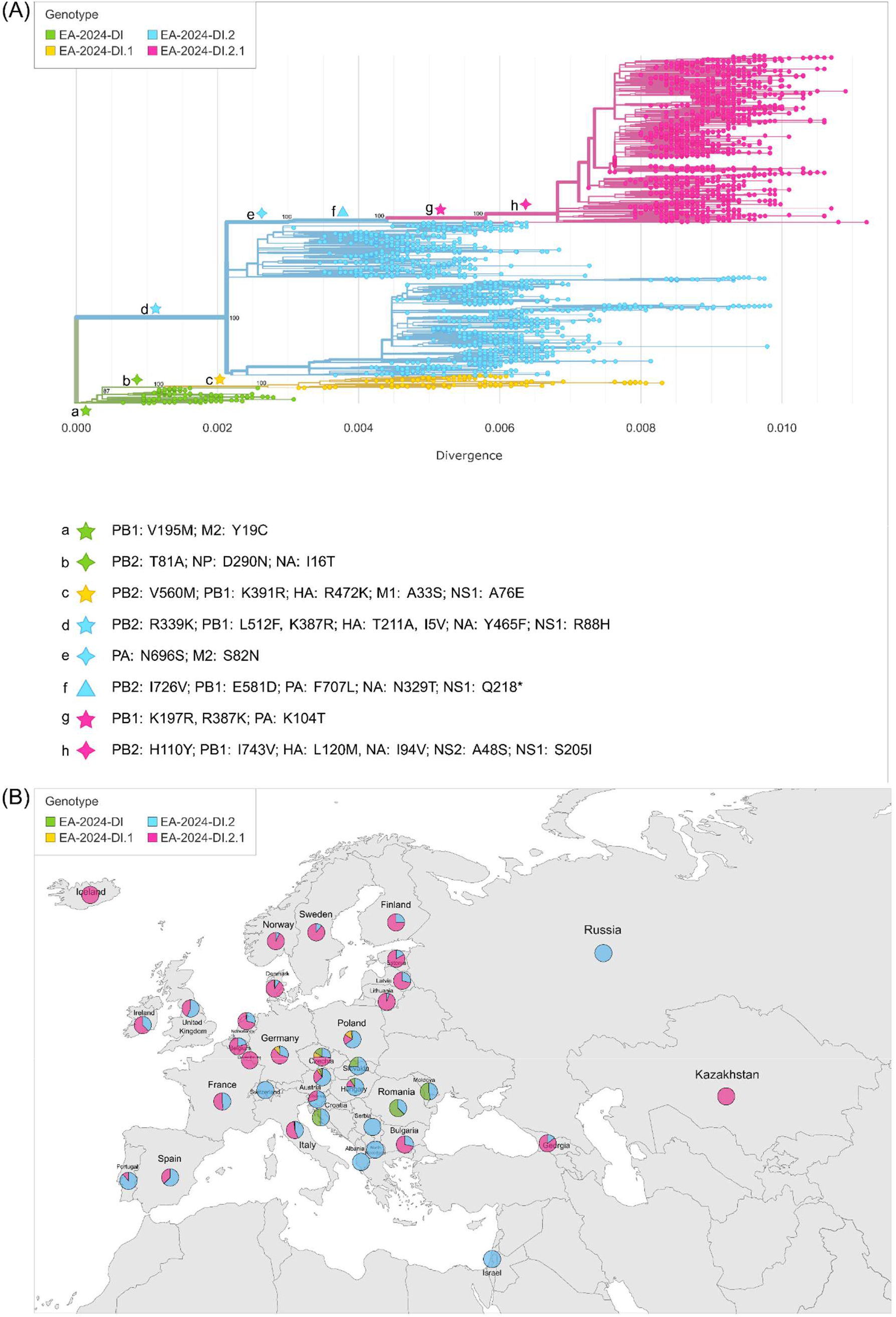
Phylogenetic tree and spatial distribution of the EA-2024-DI genotype visualized using Nextstrain. All available EA-2024-DI genotype viruses collected between December 2023 and March 2026 were used. (A) The phylogenetic tree of the complete genome of the progenitor EA-2024-DI genotype (in green) and its sub-lineages EA-2024-DI.1 (in yellow), EA-2024-DI.2 (in blue), and EA-2024-DI.2.1 (in pink). The branches defining the different sub-lineages and the amino acid substitutions acquired along each main branch are shown at bottom left. (B) Map showing the EA-2024-DI genotype and sub-lineages distribution in Europe, Russia, Kazakhstan, the Republic of Georgia and Israel.

The EA-2024-DI.1 sub-lineage was detected between June 2024 and May 2025 in eight European countries (Austria, the Czech Republic, Denmark, Germany, Italy, the Netherlands, Poland, and the United Kingdom), primarily affecting swans and domestic birds. The EA-2024-DI.2 sub-lineage began circulating in September 2024, and subsequently became widespread across Europe, being reported in 30 European countries, as well as in Israel, Russia (Krasnodar) and the Republic of Georgia (Figure 2). During the 2024–2025 epidemic wave, EA-2024-DI.2 became the predominant variant, representing 74% (N=1303/1761) of the detected and genetically characterized viruses (Figure 1). It was identified in a wide variety of wild bird species, as well as in domestic birds. Detections of EA-2024-DI.2 were also reported in mammals, including foxes, otters, cats, seals and a sheep^8^.

At the onset of the 2025–2026 epidemic wave, the EA-2024-DI.2 virus continued to circulate exclusively in Western Europe. In the rest of Europe, however, it was suddenly replaced by the EA-2024-DI.2.1 genetic drift variant, which rapidly became the predominant circulating sub-lineage across the continent (Figure 1). Phylogenetic evidence indicates that the progenitors of EA-2024-DI.2.1 were EA-2024-DI.2 viruses identified in Israel and the Republic of Georgia between late 2024 and early 2025 in domestic and wild birds. The relatively long branch separating EA-2024-DI.2.1 from these genetic progenitors suggests that the virus underwent evolutionary diversification in breeding areas within or outside Europe, where surveillance is often scarce or difficult to carry out, before its detection in September 2025 (Figure 2). Since its emergence, EA-2024-DI.2.1 has circulated extensively among wild Anseriformes with sporadic mass mortality events in swans and storks and a large mortality event in common cranes. It has also been associated with the majority of poultry outbreaks, detected in Europe since September 2025^9,10^. Interestingly, almost all characterized viruses from common cranes in European countries (Belgium, Denmark, France, Germany, Luxembourg, the Netherlands, Poland, Spain, Sweden) cluster within a monophyletic group, suggesting a single introduction event into this species followed by intra-species transmission^7^ (Figure S1 in Supplemental material). As of 31st March, 2026, the EA-2024-DI.2.1 drift variant sub-lineage had been detected in 23 European countries, as well as in the Republic of Georgia and Kazakhstan, and remains widely distributed across Europe, with ongoing detections in both wild and domestic birds.

During its evolution into different sub-lineages, the EA-2024-DI virus has acquired multiple amino acid substitutions across its genome. Figure 2 highlights the amino acid substitutions accumulated along the major branches characterizing the different sub-lineages. In particular, the EA-2024-DI.1 and EA-2024-DI.2 sub-lineages acquired, respectively, five (branch c) and seven (branch d) unique amino acid signatures compared to the EA-2024-DI progenitor. Additionally, EA-2024-DI.2.1 harbours nine further unique amino acid mutations compared to the progenitor EA-2024-DI.2 genotype (branches g and h), along with additional mutations shared with a subset of ancestral viruses within the EA-2024-DI.2 sub-lineage (branches e and f). None of these mutations has been previously described in the literature as associated with a particular phenotypic change, except for the truncated NS1 protein, 217 amino acids in length. This truncation was detected in nearly all the viruses belonging to sub-lineage EA-2024-DI.2.1 (N=1417/1420), in only 4.1% (N=55/1135) of viruses from sub-lineage EA-2024-DI.2, and in no viruses from sub-lineage EA-2024-DI.1. The role of the variation in the NS1 lengths in the virulence of avian influenza viruses in birds and mammals is highly strain-dependent; however, A(H5N8) viruses of clade 2.3.4.4b with a shorter NS1 protein have been shown to increase virulence and transmission of H5N8 in Pekin ducks^11,12^, and to be more efficient at blocking apoptosis and IFN-β response without a significant impact on virus replication in human cells^12,13^. A similar truncation is present in seasonal human A(H1N1) viruses, which contain an NS1 protein that is 219 amino acids long.

### Evolutionary and phylogeographic analyses

To enable a comparative analysis of the EA-2024-DI sub-lineages across consecutive epidemic waves, which exhibited distinct dynamics and epidemiological characteristics, both continuous and discrete phylogeographic analyses on two distinct datasets were undertaken. The first dataset (dataset 1) comprised a selection of viruses belonging to the EA-2024-DI genotype and its EA-2024-DI.1 and EA-2024-DI.2 sub-lineages, covering the 2023-2024 and 2024-2025 epidemic waves. The second dataset (dataset 2) included only viruses belonging to the EA-2024-DI.2.1 sub-lineage, which has been circulating since September 2025.

The estimate of the time to the most recent common ancestor (tMRCA) suggests that the EA-2024-DI virus originated between September and October 2023 (95% HPD: September 9– October 19, 2023). Continuous phylogeographic analysis of dataset 1 demonstrates that the EA-2024-DI virus emerged in Eastern Europe and it circulated in this region predominantly in Anseriformes and domestic birds until spring 2024, followed by progressive expansion in northern and western directions (video S1 in Supplemental material). During the 2024–2025 epidemic wave, the virus spread across Europe (Figure 3, video S1 in Supplemental material). The EA-2024-DI.1 sub-lineage was estimated to have emerged in North-Central Europe in late March 2024 (95% HPD: March 3–April 17, 2024), almost simultaneously with the EA-2024-DI.2 sub-lineage, whose ancestral node was dated to early April 2024 (95% HPD: March 8–April 17, 2024). Viruses belonging to sub-lineage EA-2024-DI.1 spread from North-eastern Europe to Western and Southern Europe (Figure 3A). In contrast, viruses of sub-lineage EA-2024-DI.2 dispersed in multiple directions from South-eastern Europe to Western and Northern European countries (Figure 3B).

**Figure 3.**
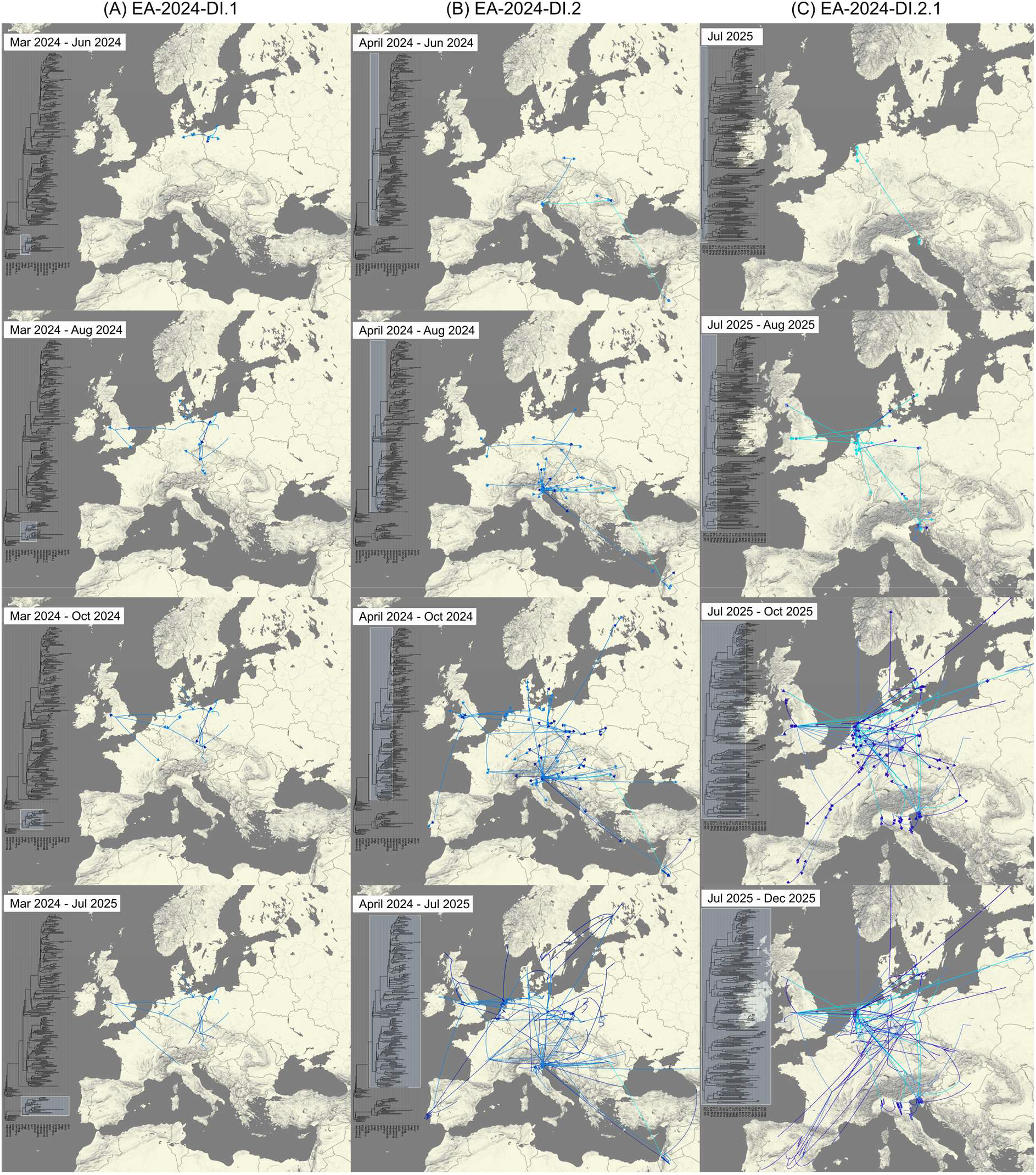
Snapshots of spatiotemporal dispersal patterns inferred from the continuous phylogeographical analysis of (A) EA-2024-DI.1 (dataset 1); (B) EA-2024-DI.2 (dataset 1); (C) EA-2024-DI.2.1 (dataset 2). Arrows are color-coded by time, using a three-level blue gradient ranging from light to dark. A time-scaled tree is displayed to the left of each snapshot. A box on the tree highlights the cluster for which transitions are displayed in columns A (EA-2024-DI.1), B (EA-2024-DI.2), and C (EA-2024-DI.2.1) on maps. The temporal bar is at the bottom of the tree.

These spatial dynamics were confirmed by the discrete phylogeographic analysis. South-eastern Europe (the yellow area in Figure 4A, comprising Albania-Bulgaria-Hungary-Moldova-Romania-Serbia, the Czech Republic-Poland-Slovakia (the dark green area), and Germany (the red area) acted as major hubs for viral dissemination. From South-eastern Europe, the virus spread northwards to the Czech Republic-Poland-Slovakia (BF=60263), westwards to Austria-Croatia-Italy-Slovenia-Switzerland (orange area, BF=60263) and to France (pink area, BF=25), and southwards to the Middle East (BF=1127). From Poland-Czech Republic-Slovakia, the virus spread further to Germany (BF=1763), France (BF=80), and Austria-Croatia-Italy-Slovenia-Switzerland (BF=80). From Germany, it reached the Scandinavian countries (blue area, BF=60263), the UK-Ireland (dark red area, BF=917), Belgium-Netherlands (light blue area, BF=699), and Spain-Portugal (light green area, BF=10). The latter regions (the Scandinavian countries, UK-Ireland, Belgium-Netherlands, Spain-Portugal) together with France and Austria-Croatia-Italy-Slovenia-Switzerland appear to have primarily served as recipients of the virus (Figure 4A, Figure S2A in Supplemental material). Markov Jumps (MJ) supported these findings, with the vast majority of jumps originating from Germany (MJ=57.6), Eastern Europe (MJ=53.3), and the Czech Republic-Poland-Slovakia (MJ=45.9) (Figure 4B). Markov rewards indicated that the virus circulated for the longest period in South-eastern Europe (Albania-Bulgaria-Hungary-Moldova-Romania-Serbia), followed by regions with comparable Markov rewards values, including the Czech Republic–Poland–Slovakia, Germany, and the UK-Ireland (Figure S3A in Supplemental material).

**Figure 4.**
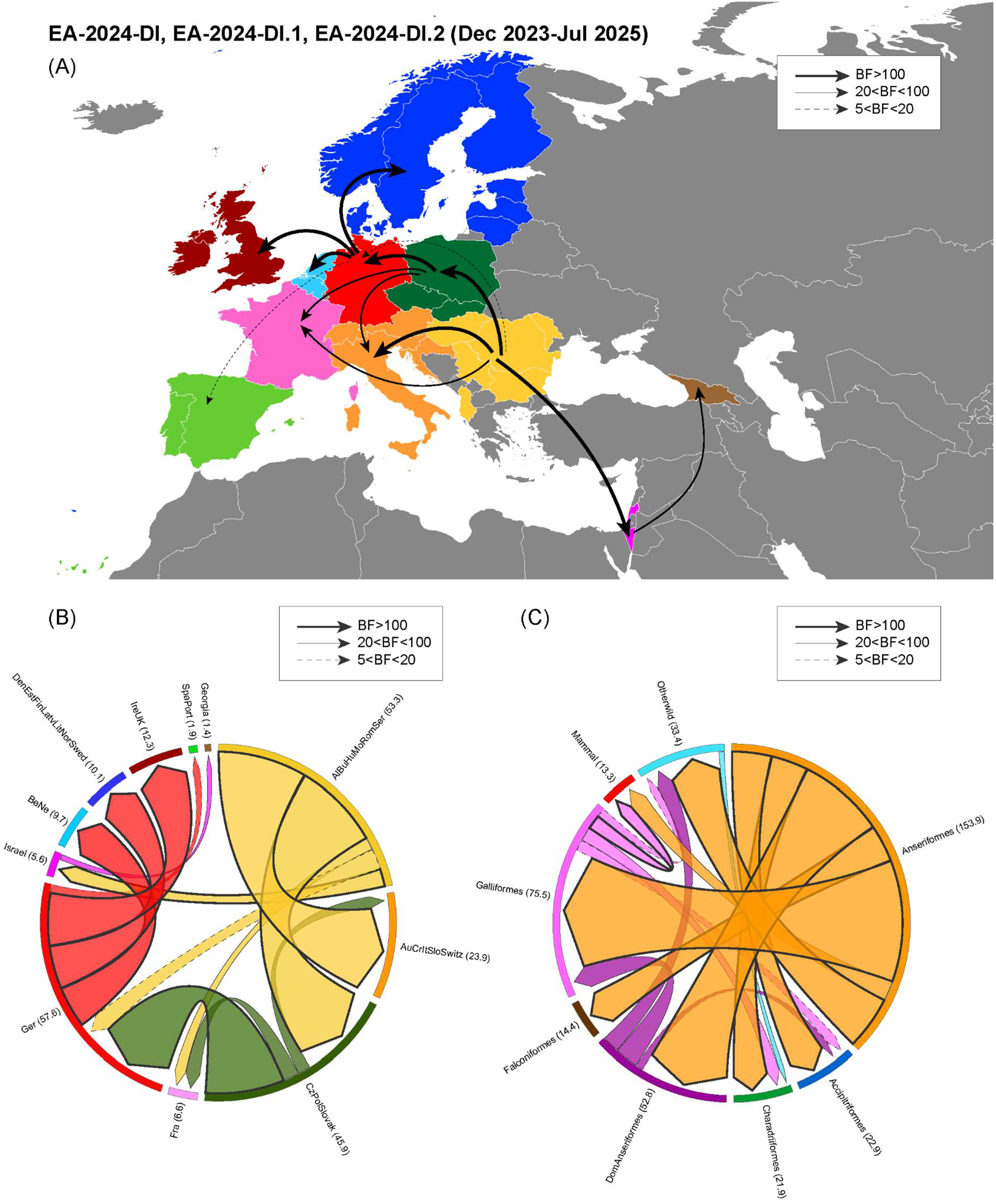
Discrete phylogeographic analysis performed in BEAST using complete genome sequences available for HPAI A(H5N1) viruses belonging to the EA-2024-DI genotype, including the EA-2024-DI.1 and EA-2024-DI.2 sub-lineages collected during the 2023-2024 and 2024-2025 epidemic waves (dataset 1). (A) Map of the discrete phylogeographic reconstruction showing transitions supported by Bayes factor (BF) > 5 and posterior probability (PP) > 0.5. (B, C) Circular migration flow plots (created using RAWGraphs, https://www.rawgraphs.io/) of EA-2024-DI genotype based on the posterior expectations of the Markov jumps for the location (B) and host (C) traits. Only transitions supported by BF > 5 and PP > 0.5 are shown. Migration flows between countries/hosts are represented by arrows starting from the location (B) or host (C) source and ending with an arrowhead at the destination location/host. The width of each arrow corresponds to the frequency of virus movement between locations/hosts, as quantified by the number of Markov jumps. The thickness of the arrow outline is proportional to the relative strength by which transitions are supported: very strong (BF > 100, thick lines), strong (20 < BF < 100, thin lines) and positive (20 < BF < 100, dashed lines). Colour code for geographic regions: Albania-Bulgaria-Hungary-Moldova-Romania-Serbia in yellow; Austria-Croatia-Italy-Slovenia-Switzerland in orange; Belgium-Netherlands in light blue; the Czech Republic-Poland-Slovakia in dark green; Denmark-Estonia-Finland-Latvia-Lithuania-Norway-Sweden in blue; France in light pink; Germany in red; Ireland-United Kingdom in dark red; Israel in pink; the Republic of Georgia in brown; Spain-Portugal in light green.

During the 2025–2026 epidemic wave (dataset 2), the dissemination pattern of the EA-2024-DI.2.1 sub-lineage appeared to differ from that observed for the EA-2024-DI, EA-2024-DI.1 and EA-2024-DI.2 sub-lineages (dataset 1). Continuous phylogeographic analysis revealed that the EA-2024-DI.2.1 sub-lineage, whose estimated tMRCA was mid July 2025 (95% HPD: June 6–August 11, 2025), spread rapidly from Central Europe to Southern, Eastern and Western countries (Figure 3C, Video S2 in the Supplemental material). Notably, the cluster of viruses associated with mass mortality events in common cranes exhibited a distinctive spread from Germany to Spain, coinciding with the western European flyway that leads the birds from their breeding area in Scandinavia to wintering grounds in France and Spain after stopovers in Germany (https://www.kraniche.de/en/crane-migration.html) (Figure 5). The discrete analysis revealed that the Belgium–Netherlands–Luxembourg (Benelux) region and Germany were the main donor areas. Virus spread from the Benelux region northwards towards the Scandinavian countries (BF=65541) and the UK (BF=65541), eastwards to Germany (BF=65541) and the Czech Republic-Poland (BF=1865), and southwards to the region comprising Austria and Italy (BF=1358). Dissemination from Germany predominantly occurred westwards to the Benelux region (BF=129), France (BF=7275) and Spain (BF=65541), possibly in association with the main migration route of common cranes within Europe (Western European Flyway) (Figure 6A, Figure S4A in supplemental material)^14^. MJ analyses supported these findings (Figure 6B), and Markov rewards indicated that the virus spent the longest period in the Benelux region, followed by Germany and in the region comprising Austria and Italy (Figure S3C in Supplemental material).

**Figure 5.**
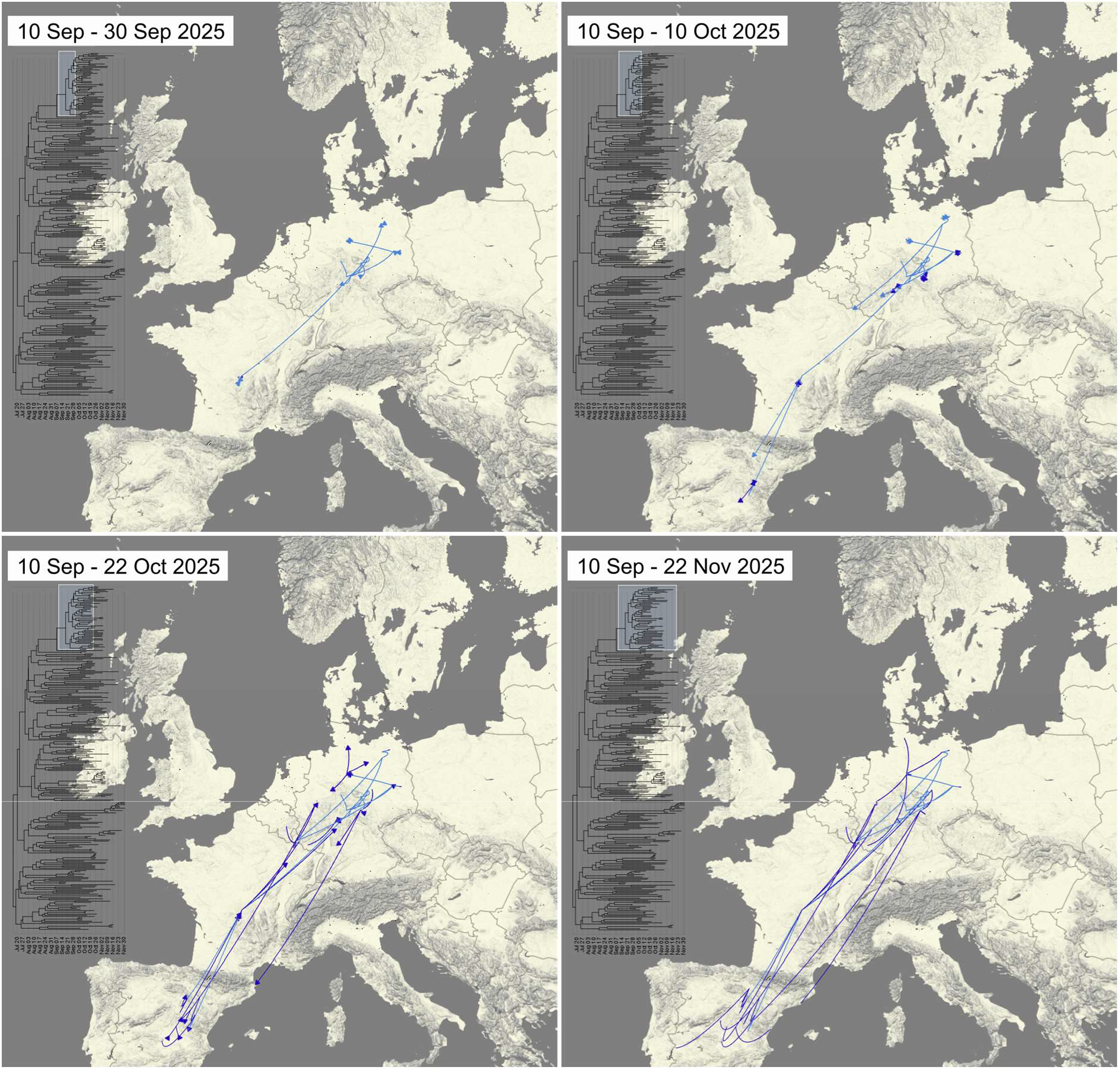
Snapshots of spatiotemporal dispersal patterns inferred from the continuous phylogeographical analysis of the EA-2024-DI.2.1 viruses (dataset 2), belonging to the common crane cluster. Arrows are colour-coded by time, using a two-level blue gradient ranging from light to dark. A time-scaled tree is displayed to the left of each snapshot. A box on the tree highlights the common crane cluster for which transitions are displayed on maps. The temporal bar is at the bottom of the tree.

**Figure 6.**
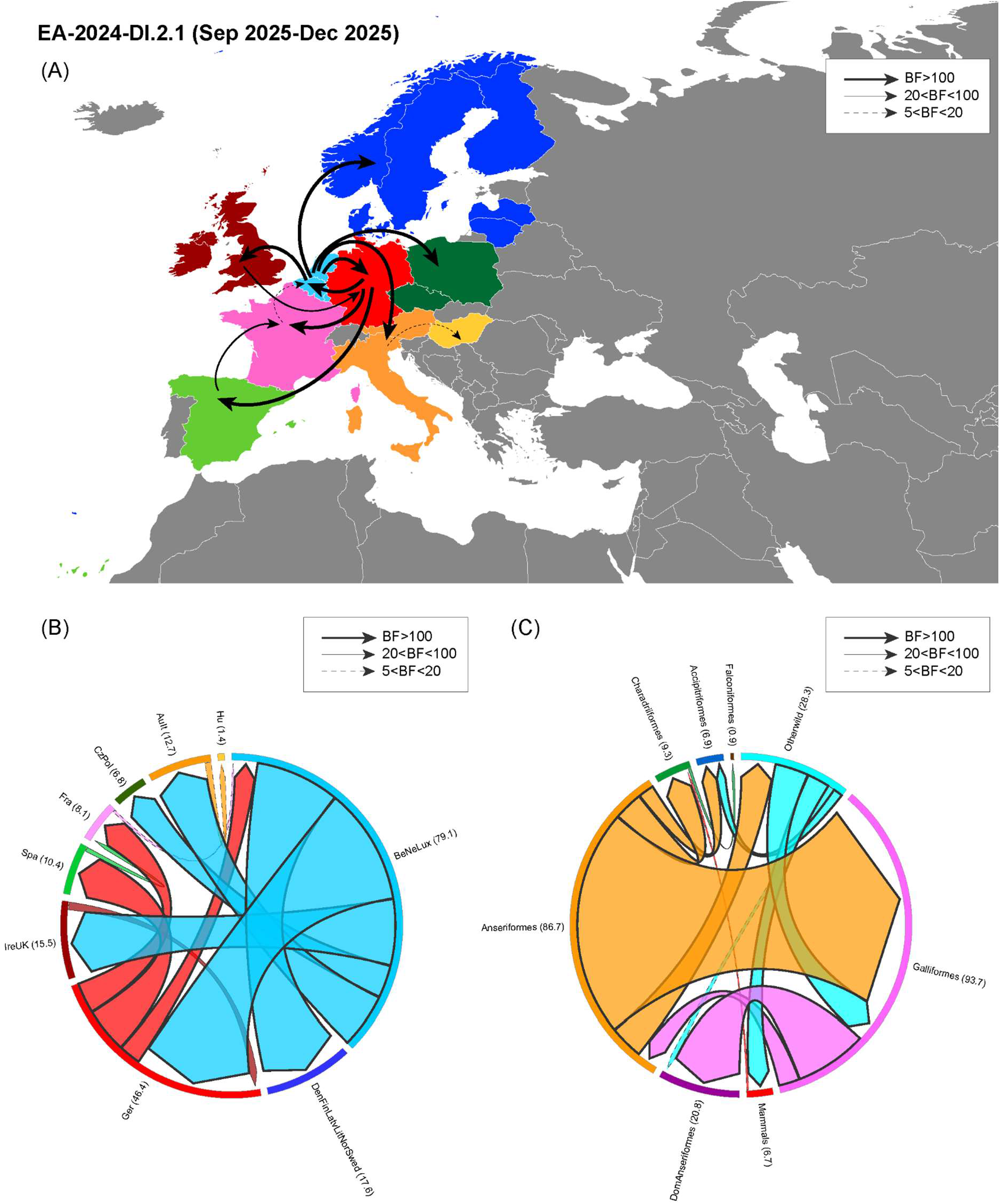
Discrete phylogeographic analysis performed in BEAST using complete genome sequences available for HPAI A(H5N1) viruses belonging to the EA-2024-DI.2.1 sub-lineage collected between September and December 2025 (dataset 2). (A) Map of the discrete phylogeographic reconstruction showing transitions supported by Bayes factor (BF) > 5 and posterior probability (PP) > 0.5. (B, C) Circular migration flow plots (created using RAWGraphs, https://www.rawgraphs.io/) of EA-2024-DI genotype based on the posterior expectations of the Markov jumps for the location (B) and host (C) traits. Only transitions supported by BF > 5 and PP > 0.5 are shown. Migration flows between countries/hosts are represented by arrows starting from the location (B) or host (C) source and ending with an arrowhead at the destination location/host. The width of each arrow corresponds to the frequency of virus movement between locations/hosts, as quantified by the number of Markov jumps. The thickness of the arrow outline is proportional to the relative strength by which transitions are supported: very strong (BF > 100, thick lines), strong (20 < BF < 100, thin lines) and positive (20 < BF < 100, dashed lines). Colour code for geographic regions: Hungary in yellow; Austria-Italy in orange; Belgium-Netherlands-Luxembourg in light blue; the Czech Republic-Poland in dark green; Denmark-Finland-Latvia-Lithuania-Norway-Sweden in blue; France in light pink; Germany in red; Ireland-United Kingdom in dark red; Spain in light green.

### Host contribution

To disentangle the contribution of different wild and domestic host species to the dissemination of the different EA-2024-DI sub-lineages, we performed discrete phylodynamic analyses of the two datasets, treating different host groups (described in the Material and Methods and in Supplementary Table S1) as a discrete trait. For dataset 1 (2023-2025, EA-2024-DI, EA-2024-DI.1 and EA-2024-DI.2), Markov rewards indicated that the virus persisted longest in wild Anseriformes (Figure S3B in Supplemental material), likely contributing to its transmission (MJ=153.9) to other host categories (Falconiformes and Galliformes, BF=40745; Charadriiformes and other wild bird species, BF=10181; domestic Anseriformes, BF=2257; Accipitriformes, BF=422; mammals, BF=29) (Figure 4C). Poultry predominantly acted as a sink host, with wild Anseriformes representing the main source of infection for the virus. However, several transmission events from poultry to wildlife were also detected, including Galliformes to Charadriiformes (BF=93), Galliformes to other wild bird species (BF=16), Galliformes to Accipitriformes (BF=11), domestic Anseriformes to other wild bird species (BF=63) and domestic Anseriformes to Acciptriformes (BF=14). Mammals appeared to be primarily infected following assumed interactions with Anseriformes (BF=29) and Galliformes (BF=104) (Figure 4C, Figure S2B in Supplemental material).

The hosts involved in disseminating the EA-2024-DI.2.1 sub-lineage (dataset 2) were mainly Anseriformes (MJ=86.7) and “other wild birds” (MJ=28.3). Anseriformes contributed to the virus spread to Charadriiformes (BF=56512), Galliformes (BF=56512), other wild bird species (BF=56512) and to Accipitriformes (BF=158). “Other wild birds”, mainly represented by Gruiformes in dataset 2, particularly the common crane (*Grus grus*), also acted as an important source of virus for Galliformes (BF=56512), mammals (BF=56512) and Accipitriformes (BF=2087) (Figure 6C, Figure S4B in supplemental material). The pivotal role of “other wild birds” as a source of virus for other wild and domestic bird species, as well as mammals, was not observed in dataset 1.

### Mutations in the PB2 protein associated with virus adaptation in mammals

Mutations in the PB2 protein associated with virus adaptation in mammals (E627K/V, D701N and K526R)^15^ were identified in 72 viruses of the EA-2024-DI genotype: 51 (1.7%) were collected from avian species (N=2951) whereas 21 (50%) were detected in mammalian species (N=41). Based on the available data, the frequency of PB2 mutations in EA-2024-DI, EA-2024-DI.1 and EA-2024-DI.2 sub-lineages was approximately 20% (4/21) in mammals and 3.2% (44/1375) in avian species during the 2024–2025 epidemiological year, with the majority of viruses containing these mutations belonging to the sub-lineage EA-2024-DI.2. In contrast, the proportion of PB2 mutations in mammals increased markedly in 2025–2026, reaching up to 80% (16/20) (Figure 7), while in avian species it dropped to 0.2% (3/1399). The high frequency of viruses with a zoonotic PB2 mutation detected in birds during the 2024-2025 epidemic was due to the clusters of viruses containing PB2 molecular markers of mammalian adaptation identified in both wild and domestic birds (Figure S5 in Supplemental material). This suggests that viruses carrying these mutations are capable of efficient transmission in avian populations. By contrast, no clusters of viruses containing PB2 mutations were detected in birds during the 2025-2026 epidemic wave.

**Figure 7.**
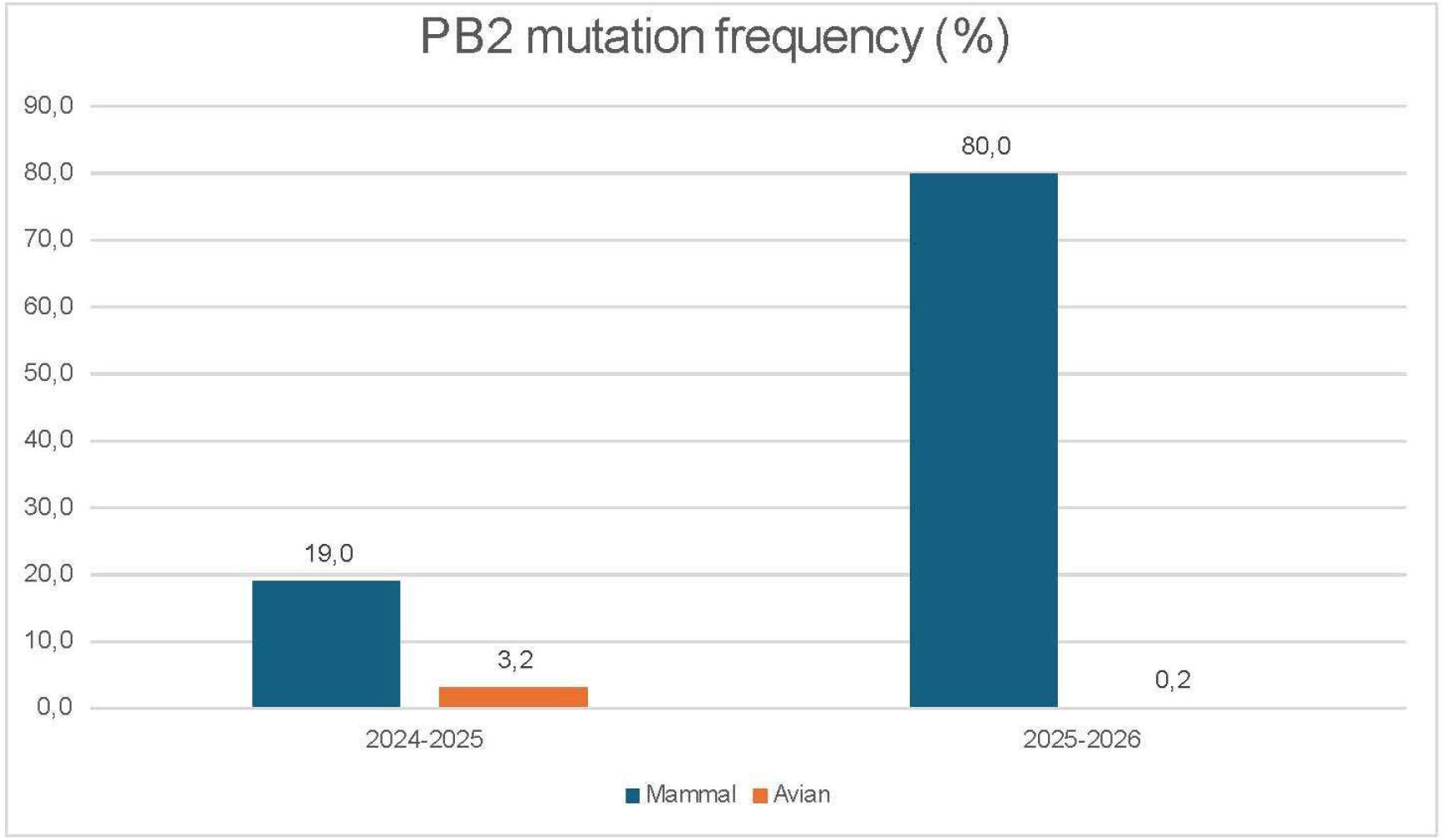
Frequency of mutations in the PB2 protein associated with virus adaptation in mammals. The graph includes data from mammals and avian species collected from October 2024 to the end of March 2026.

## Discussion

Since the introduction of the HPAI A(H5) of clade 2.3.4.4b in 2020, European epidemic waves have been characterized by the emergence of new genotypes with an ever-increasing ability to infect a broad range of wild birds^4^. From 2020 to 2023, a rapid turnover of genotypes was observed, most of which became extinct after one or two epidemic waves, whilst others continued to circulate persistently at low frequencies in well-defined ecological niches (e.g., genotypes EA-2021-I, EA-2024-DT, EA-2022-BB andEA-2024-DA)^4,6,16^. Since October 2024, for the first time in Europe, a single genotype, EA-2024-DI, has driven two successive epidemic waves, 2024-2025 and 2025-2026, but with a completely different outcome. An unprecedented number of cases in wild birds was recorded during the 2025-2026 epidemic wave, representing a six-fold rise compared to the equivalent period (1 October-31 March) of the 2024-2025 wave (https://eurlaidata.izsvenezie.it/). What drove this exceptionally high level of viral circulation in the wild avian population? This study reveals that, during the 2024-2025 epidemic, two sub-lineages, EA-2024-DI.1 and EA-2024-DI.2 were co-circulating. These variants evolved from the EA-2024-DI genotype that emerged in Europe in late 2023. The current 2025-2026 epidemic, however, has been caused by a newly emerged drift variant of EA-2024-DI.2, namely EA-2024-DI.2.1. Our results indicate that wild Anseriformes were the principal drivers of viral dissemination across most host categories for EA-2024-DI sub-lineages. This central role of wild Anseriformes is consistent with their known ecological characteristics, such as high abundance, gregarious behaviour, frequent congregations at wetlands, and long-distance migration. A distinctive feature of EA-2024-DI.2.1 is the involvement of common cranes, alongside Anseriformes, as important drivers to viral spread. This may partially explain the initial peak of infection observed in wild birds in October-November 2025^17^. The inferred dissemination route of the viruses collected from the common cranes aligns with the Western European Flyway, which connects common cranes’ breeding sites in Scandinavia, Germany, and Poland to wintering grounds in Spain and Portugal. (https://www.kraniche.de/en/crane-migration.html). The incursion of the virus into this species during migration, along with the unprecedented mass mortality events observed, may be one factor contributing to the rapid dissemination of the EA-2024-DI.2.1 viruses South-westward. The clear clustering of viruses collected from common cranes suggests a single introduction event in this species, followed by intra-species transmission and occasional spillover events to other wild and domestic bird species, as well as wild mammals. This cluster has been identified exclusively in countries through which common cranes migrate, with the last detection dating back to the end of January 2026, which may be indicative of a localized and self-contained event. Thus, other factors may have contributed to the differing viral dissemination during the two epidemic waves.

Anseriformes have also been identified as the main donor of the genotype EA-2024-DI for domestic birds. For the EA-2024-DI.2.1 sub-lineage, the category ‘Other wild birds’, which mainly includes common cranes, has been identified as a second major source of virus transmission to domestic species. However, despite the various sources of the virus for domestic birds associated with EA-2024-DI.2.1, the overall number of outbreaks in poultry reported since September 2025 is comparable to that of the previous season^6^. This observation may be explained by the lower level of farm-to-farm transmission reported during 2025–2026 epidemic compared to the previous wave^6,18^.

These data suggest that the EA-2024-DI.2.1 drift variant represents a novel introduction of a virus into Europe that occurred during the 2025 fall migration of wild birds. This is indicated by the number of amino acid substitutions (N=9) and the long branch separating EA-2024-DI.2.1 from its progenitor viruses EA-2024-DI.2, collected in Israel and the Republic of Georgia during the winter of 2024-2025. This gap in the data prevents us from reconstructing the complete emergence and dissemination of this novel sub-lineage. However, the spatial reconstruction identifies the North Sea coast of the Netherlands as the most likely entry point of EA-2024-DI.2.1 into Western Europe. This area lies along the East Atlantic flyway, which is used by several duck species during their fall migration from Siberia to Europe^19^. It has also been identified as a major European hotspot for the introduction of HPAI H5 viruses from breeding grounds in northern Russia during previous epidemic waves^20,21^. The new incursion with infected migratory birds during the 2025 fall migration may have facilitated the rapid dissemination of this variant in Europe. Above-average snow cover and lower-than-average temperatures reported in September 2025 in Northen and Northeastern Russia (https://climateimpactcompany.com/daily-feature-expanding-early-season-snow-cover-in-russia-2/), were not observed in 2024. These conditions may have triggered the early arrival of infected migratory birds and their contact with other European species that had not previously been exposed to avian influenza viruses, including juvenile individuals, as well as with short-to medium-distance migratory species within Europe, such as common cranes. This species on the Western European flyway was rarely affected during previous epidemics, facilitating increased viral spread^17^.

These data suggest that the EA-2024-DI genotype emerged in Europe in late 2023. While we cannot rule out the possibility of new introductions of EA-2024-DI.1 and EA-2024-DI.2 in Europe during the fall migration of 2024, the absence of any progenitor viruses being detected outside Europe is consistent with the evolutionary pathway occurring within Europe. Moreover, the continuous phylogeographic analysis indicates a persistent circulation of this genotype in Eastern Europe throughout the 2023-2024 epidemic. Both sub-lineages likely emerged in late March and early April 2024, respectively, based on the estimated tMRCAs, and began spreading Westward at the end of summer 2024. This occurred immediately before the start of the 2024-2025 epidemic and coincided with an increase in the wild Anseriformes population moving into European wintering areas.

Other factors associated with the intrinsic characteristics of the different sub-lineages, may have influenced the differing courses of the two epidemics. A notable feature of EA-2024-DI.2.1 is its truncated NS1 protein, which may contribute to enhanced viral transmission. This truncation was also present in the 2016 A(H5N8) viruses of clade 2.3.4.4b, where it was associated with enhanced viral replication and bird-to-bird transmission in ducks, as well as increased virulence in mice^11^. However, the role of this truncation in host adaptation, pathogenicity and transmission remains unclear, and further studies are needed to understand its effect on this specific strain.

During the 2025-2026 epidemic (up to February 2026), an increase in the number of A(H5N1) cases in mammals was observed in Europe (over 40 cases) compared to the same time period (October-February) in 2024-2025 (approximately 10 cases). During both epidemics, nearly all the viruses collected from mammals belonged to the EA-2024-DI genotype^6,7,18,22^. The increase in mammalian infections is likely associated with the higher levels of virus circulation in birds during the 2025-2026 epidemic and with a high environmental contamination from the mass mortality events involving common cranes. These events emerged as a major source of virus exposure for mammals at the beginning of the 2025-2026 epidemic wave (October-November 2025), although it cannot be ruled out that surveillance was stepped up following the die-off of common cranes. Of note, the genetic data clearly demonstrate a significant increase in the frequency of the PB2 molecular markers of mammalian adaptation in the EA-2024-DI viruses collected from mammals, when compared to previous seasons. The frequency of detection for these markers increased from approximately 20% during the 2024-2025 epidemic to 80% during the 2025-2026 epidemic. Previous studies have demonstrated that these mutations can be rapidly acquired by viruses of clade 2.3.4.4b during their replication in naturally and experimentally infected mammalian hosts. This enhances replication, transmission of these viruses and disease progression rates in experimentally infected species^23–26^. Several factors could account for this difference in emergence of key mutations. One possibility is that viruses of the EA-2024-DI.2.1 sub-lineage are predisposed to acquiring such mutations through a combination of viral and host factors. Further studies are required to understand the implications of increased mutability of these viruses in different settings. Despite the high propensity of clade 2.3.4.4b viruses to undergo reassortment events, this study demonstrates that a single, high-fitness genotype can become established in bird populations and maintain its genomic composition over time, evolving into sub-lineages. Overall, these findings indicate that genetic composition is not the only factor shaping the outcome of an outbreak. Further studies are needed to better understand the impact of environmental and ecological factors on viral spread and variability in transmission rates and virulence among sub-lineages in avian and mammalian hosts. This study has several limitations, primarily due to heterogeneity in surveillance efforts, particularly active surveillance, and sequencing capacity across European countries. Additionally, bias due to the scarcity of data from viruses collected outside Europe cannot be excluded. Despite these limitations, this study provides the first comprehensive and up-to-date overview of the evolutionary trends of the EA-2024-DI genotype of clade 2.3.4.4b, which is responsible for the most devastating epidemic ever recorded among wild birds in mainland Europe. Our findings indicate that changes in affected wild bird species and in viral transmission patterns can influence how epidemic develop and the impact of the virus on mammals, providing crucial information for disease risk management in a One-Health context.

## Methods

### Nextstrain evolutionary analyses

The geographic diffusion, evolution and accumulation of mutations in the separate sub-lineages of the EA-2024-DI genotype were assessed using Nextstrain (https://nextstrain.org)^27^. For this analysis, the complete genome sequences of all the EA-2024-DI viruses (N=2996) collected up to March 31, 2026 were used (EA-2024-DI, N = 131; EA-2024-DI.1, N = 110; EA-2024-DI.2, N = 1335; EA-2024-DI.2.1, N = 1420), in order to provide the most up-to-date analysis. The dataset includes viruses from 31 European countries, as well as from Russia, Kazakhstan, The Republic of Georgia, and Israel.

Sequences were genotyped, aligned, and concatenated as described above (Datasets for the Bayesian phylodynamic analyses). Phylogenetic and evolutionary analyses of the EA-2024-DI genotype were performed using Nextstrain, integrating complete genome sequences with associated metadata (sampling date, country of origin and sub-lineage). The Nextstrain time-resolved tree was visualized using Auspice v.28.0.1. The time-calibrated phylogeny was reconstructed and annotated with geographic information. Nucleotide and amino acid substitutions were mapped onto the tree.

### Datasets for the Bayesian phylodynamic analyses

All available whole-genome sequences of the EA-2024-DI genotype as of December 10, 2025 (n=1,798; including N = 169 DI, N = 71 DI.1, N = 1,179 DI.2, and N = 379 DI.2.1) were collected. These genome sequences were obtained either through direct sequencing at the European Union Reference Laboratory (EURL) for avian influenza and Newcastle disease, through sequencing at the respective National Reference Laboratories for avian influenza, or by retrieving them from the GISAID database (http://www.gisaid.org, accessed on 10/12/2025). Genotypes were identified by analysing the topology of phylogenetic trees for each gene segment as described by Fusaro et al., 2024^4^ and by using the Genin2 tool version 2.1.6 (https://github.com/izsvenezie-virology/genin2). AIV genome sequences were concatenated (PB2-PB1-PA-HA-NP-NA-MP-NS) and aligned using MAFFT v7^28^. They were then tested to exclude any recombinants with RDP5^29^ and analysed using a maximum likelihood (ML) approach^30^ in IQ-TREE v3.0.1, with the substitution model GTR + F + I + R3, automatically selected according to the Bayesian Information Criterion using ModelFinder^31^. The statistical support for each node was assessed via ultrafast bootstrap approximation approach with 1,000 replicates^32^.

Two different datasets were generated. Dataset 1 included viruses collected during the 2023-2024 and 2024-2025 epidemic waves (genotypes EA-2024-DI, EA-2024-DI.1 and EA-2024-DI.2). It comprised 1,419 viral sequences collected from 29 European countries, the Republic of Georgia and Israel between December 15, 2023 and July 31, 2025. Dataset 2 consisted of 379 viral sequences of the drift variant sub-genotype EA-2024-DI.2.1 collected between September and December 2025 from 19 European countries, the Republic of Georgia and Israel. Eleven discrete regions were defined:

Albania-Bulgaria-Hungary-Moldova-Romania-Serbia; Austria-Croatia-Italy-Slovenia-Switzerland; Belgium-Netherlands-Luxembourg; the Czech Republic-Poland-Slovakia; Denmark-Estonia-Finland-Latvia-Lithuania-Norway-Sweden; France; Germany; Ireland-the United Kingdom; Israel; the Republic of Georgia; Spain-Portugal. Eight host categories were defined: Mammals (wild and domestic), Accipitriformes, Anseriformes, Charadriiformes, Domestic-Anseriformes, Falconiformes, Galliformes, Other wild birds (including Casuariiformes, Ciconiiformes, Columbiformes, Coraciiformes, Gruiformes, Passeriformes, Pelecaniformes, Podicipediformes, Rheiformes, Strigiformes, Suliformes) (Supplementary Table S1).

TempEst v1.5.3 (https://tree.bio.ed.ac.uk/software/tempest/) was used to assess the temporal signal of both datasets and to remove outliers^33^. Subsampling of each dataset was undertaken using SAMPI (https://github.com/jlcherry/SAMPI) to ensure balanced representation by location, host and sampling time. This strategy minimizes sampling bias and ensures temporal, host and geographic representativeness for robust evolutionary analysis. The final sub-sampled datasets (Supplementary Table S1, Supplementary Table S2) consisted of n = 273 (dataset 1) and n = 292 (dataset 2) complete genome sequences. Metadata are available in Supplementary Table S1.

### Evolutionary and phylogeographic analyses

To investigate the emergence, host contribution, and spread dynamics of the EA-2024-DI genotype and its drift variants, we applied a Bayesian framework based on Markov Chain Monte Carlo (MCMC) methods implemented in BEAST v1.10.4 and the BEAGLE library^34^. Multiple clock models and tree priors were compared and the best-fitting model was selected using the path sampling and stepping-stone sampling marginal likelihood estimators as implemented in BEAST^35^. Analyses were conducted using an uncorrelated lognormal relaxed molecular clock, the SRD06 nucleotide substitution model, and a Coalescent GMRF Bayesian Skyride as the tree prior. We ran the analyses until all relevant effective sample sizes reached at least 200, as assessed in Tracer v1.7.2^36^ and checked if independent replicate analyses converged to the same posterior distribution. Maximum clade credibility (MCC) trees were summarized using TreeAnnotator v10.5.0, after removing 10% of the chain as burn-in, and visualized with FigTree v1.4.4 (http://tree.bio.ed.ac.uk/software/figtree/).

Discrete phylogeographic analyses were performed using location and host traits, using the categories described above. An asymmetric transition model was assumed, and Bayesian stochastic search variable selection was incorporated (Lemey et al., 2009). Bayes factor (BF) support for individual transitions between discrete locations and hosts was assessed using Spread D3 v0.9.6^37^, with BF support interpreted as positive (5 ≤ BF < 20), strong (20 ≤ BF < 100), and very strong (BF ≥ 100)^38^. Markov jump counts were used to estimate the number of viral movements along the phylogenetic branches, and Markov rewards quantified the time the virus spent in each host and geographic region^39^.

Continuous phylogeographic analyses were performed on the same datasets (datasets 1 and 2), with location data provided as latitude and longitude coordinates. A gamma-relaxed random walk (RRW) diffusion model was applied, allowing dispersal rates to vary across branches of the phylogenetic tree. The MCMC was run for 500 million and 1 billion iterations for dataset 1 and dataset 2, respectively.

EvoLaps version 3 (https://www.evolaps.org/evolaps3.22/index.html) was used to visualize continuous phylogeographic reconstructions^40^.

### Molecular analysis

Molecular markers associated with zoonotic potential were identified in viral proteins using the FluMut tool (FluMut version 0.6.4, FluMut database version 6.5) (https://izsvenezie-virology.github.io/FluMut/)^41^.

## Acknowledgements

The authors wish to thank Francesca Ellero for her careful revision of the manuscript and all the colleagues in the Viral Genomics and Transcriptomics Laboratory and Experimental Virology Laboratory at the IZSVe for their support in the diagnosis and characterization of avian influenza viruses. We gratefully acknowledge also the authors, originating and submitting laboratories of the sequences from GISAID’s EpiFlu^™^ Database on which this research is based (https://doi.org/10.55876/gis8.260521qn).

