## Supplementary material for "Two epidemics, one genotype, different outcomes: evolutionary changes of Avian Influenza H5N1, genotype EA-2024-DI": Supplemental material.docx

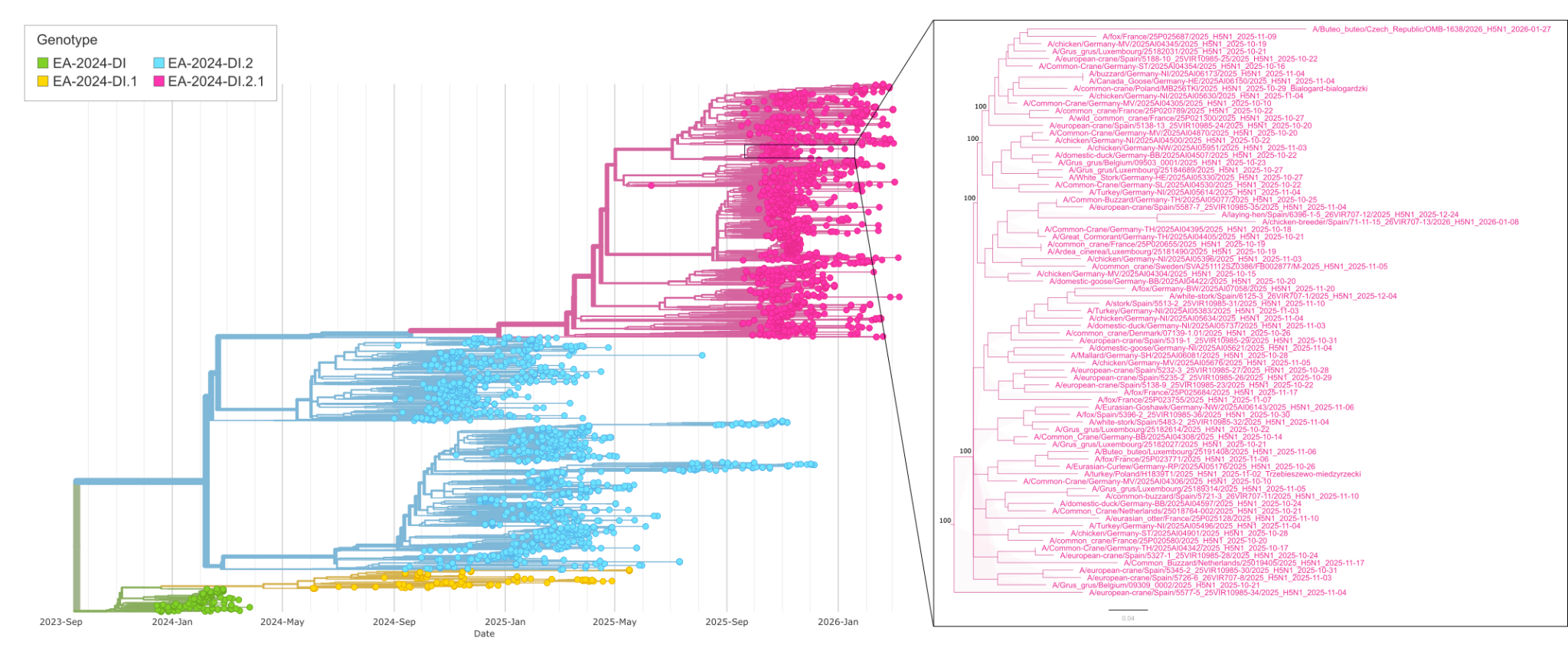

**Figure S1.** Phylogenetic tree of the EA-2024-DI genotype visualized using Nextstrain. In the right panel, a magnified view of the common crane cluster is shown.

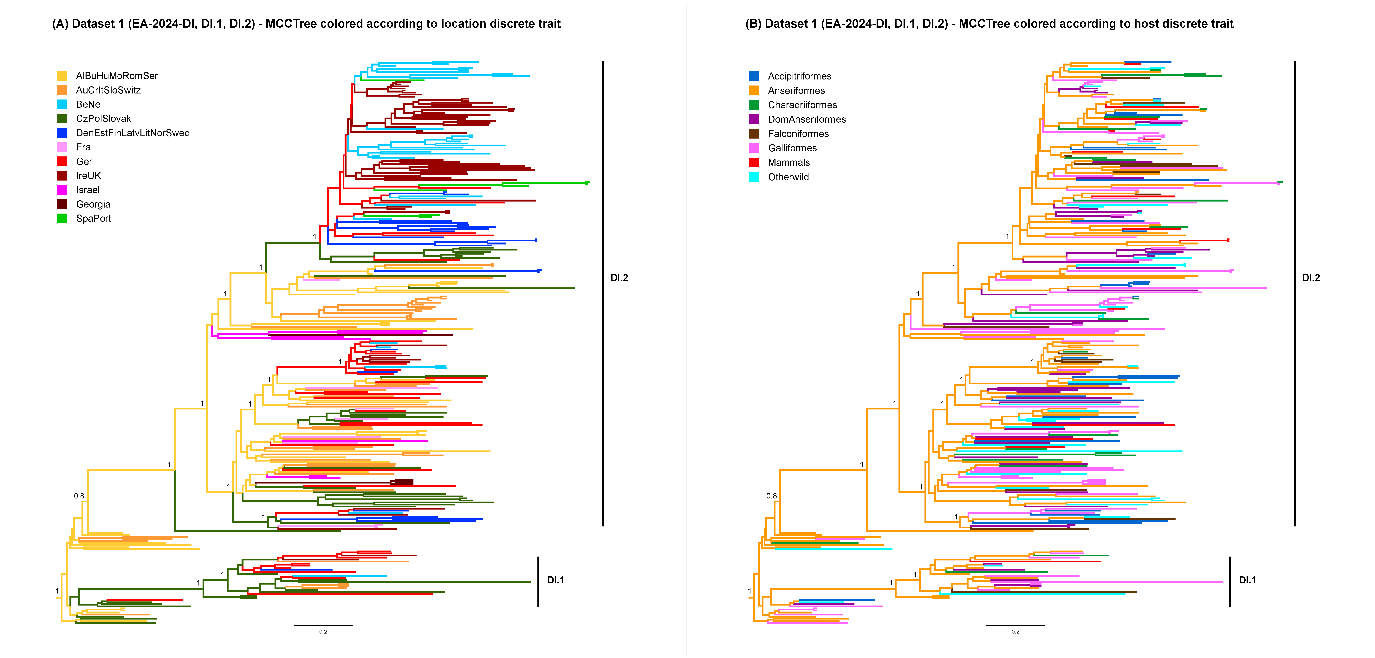

**Figure S2**. Maximum Clade Credibility tree of the EA-2024-DI genotype and its EA-2024-DI.1 and EA-2024-DI.2 sub-lineages (dataset 1), coloured according to (A) the location discrete trait and (B) the host discrete trait.

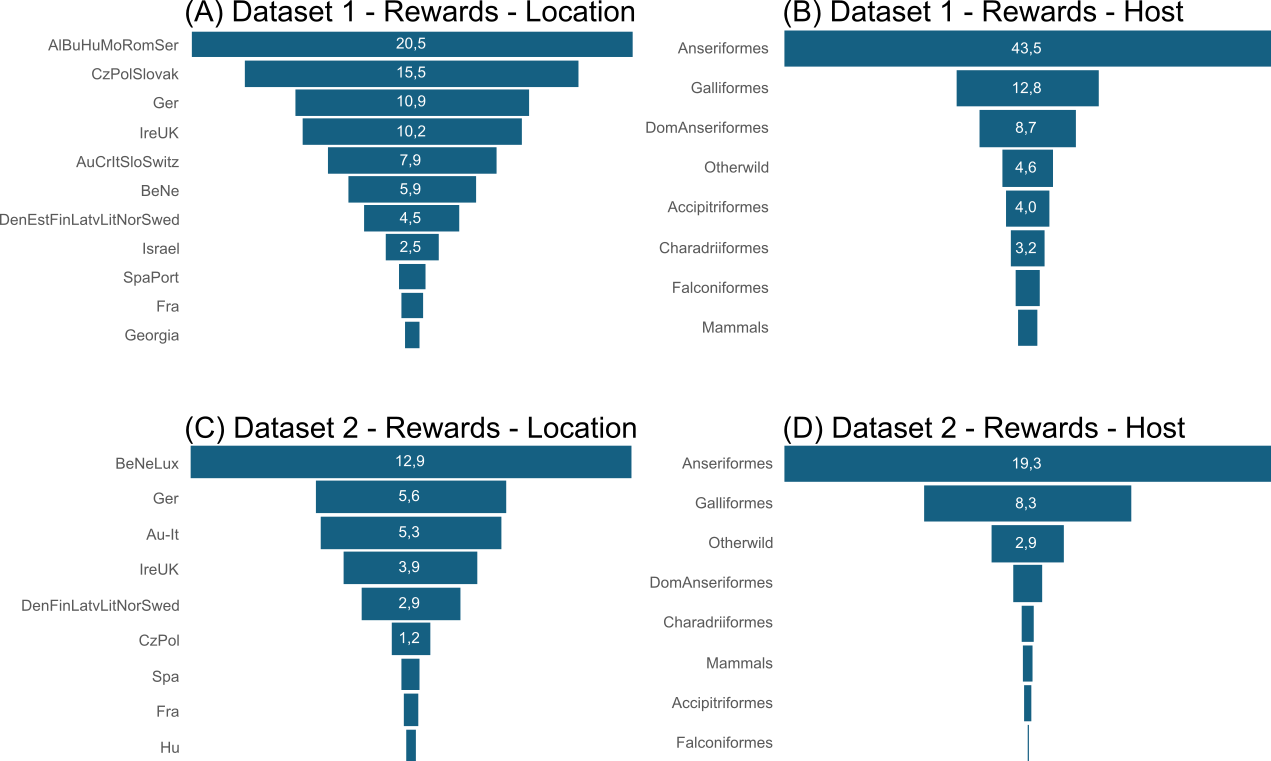

**Figure S3.** Markov rewards for (A) location trait in dataset 1; (B) host trait in dataset 1; (C) location trait in dataset 2; (D) host trait in dataset 2. Dataset 1 consists of viruses belonging to the EA-2024-DI genotype and its EA-2024-DI.1 and EA-2024-DI.2 sub-lineages; dataset 2 consists of viruses belonging to the EA-2024-DI.2.1 sub-lineage.

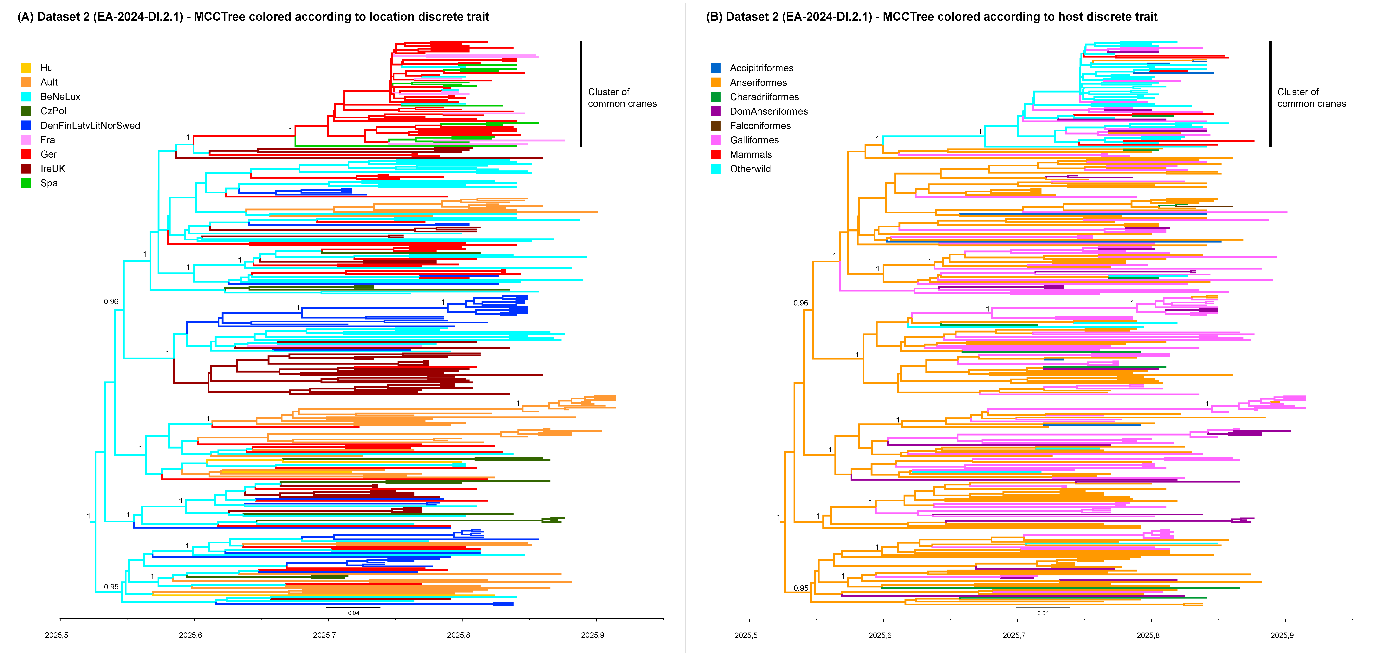

**Figure S4**. Maximum Clade Credibility tree of the EA-2024-DI.2.1 sub-lineage (dataset 2), coloured according to (A) the location discrete trait and (B) the host discrete trait.

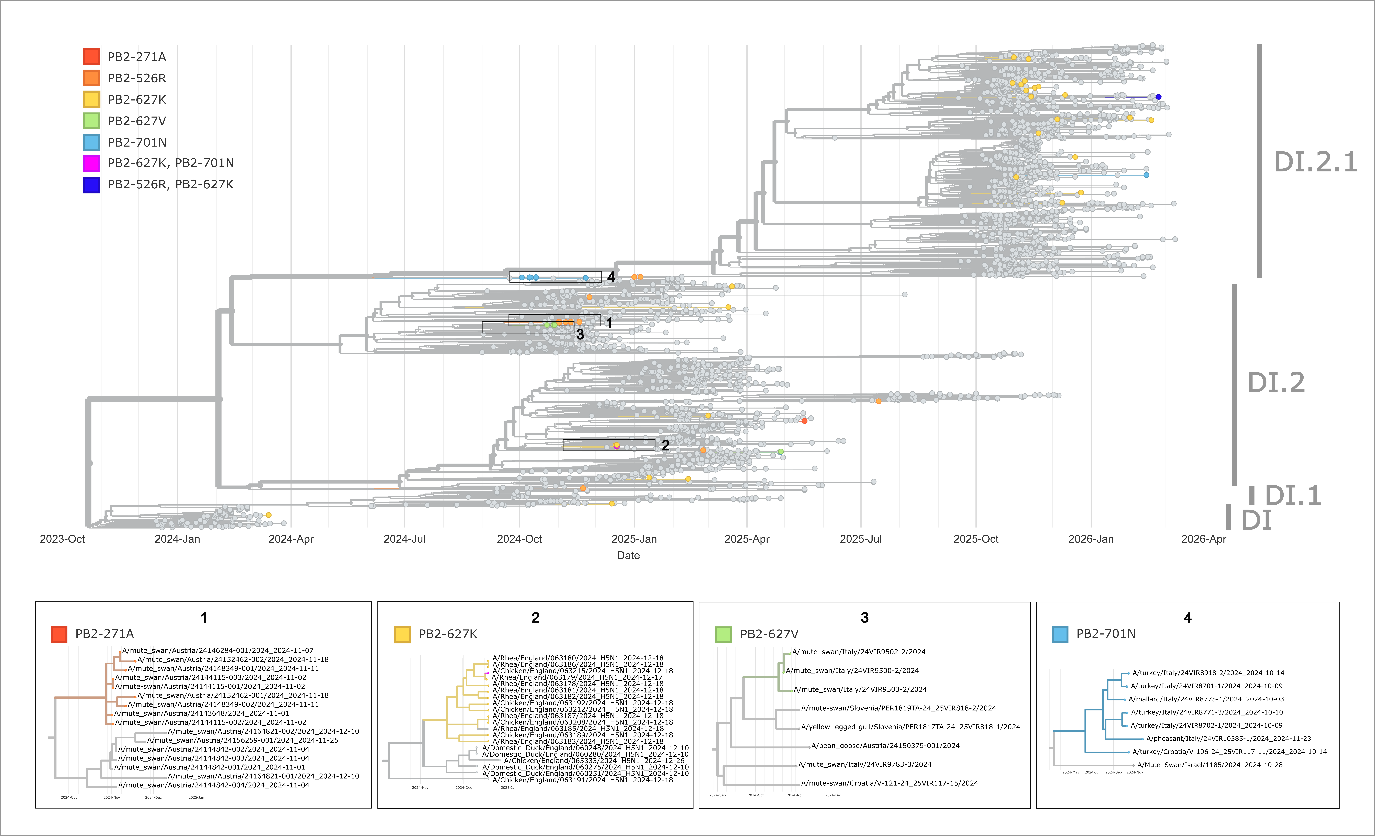

**Figure S5.** EA-2024-DI viruses containing PB2 molecular markers of mammalian adaptation are highlighted in the phylogenetic tree. Clusters of viruses bearing the same mutation are shown in the four boxes at the bottom.
