## Supplementary material for "Two epidemics, one genotype, different outcomes: evolutionary changes of Avian Influenza H5N1, genotype EA-2024-DI": Table S2: SupplementaryTableS2_gisaid_supplemental_table_epi_set_260521qn.pdf

### Supplementary Appendix

All genome sequences and associated metadata supporting the findings of this study can be accessed through the persistent digital object identifier https://doi.org/10.55876/gis8.260521qn

In addition to the minted DOI, GISAID also communicates the aggregation of GISAID accession numbers (EPI_ISL_IDs) through the corresponding EPI_SET_260521qn identifier to facilitate both, the acknowledgment of all data contributors and the direct retrieval of the underlying data from GISAID used in this study.

##### Influenza Virus Data Summary

| GISAID Identifier | Digital Object Identifier | Number of individual viruses | Data Collection range | Number of countries/territories |
|---|---|---|---|---|
| EPI_SET_260521qn | https://doi.org/10.55876/gis8.260521qn | 565 | 2023-12-15 to 2025-12-01 | 29 |

**PLEASE NOTE!**

The following data are placed under a temporary publishing embargo:

20267041 until 03-Jun-2026 00:00 UTC

These data cannot be used in a publication of any kind until the temporary publishing embargo has expired, unless, a prior written consent of the Data Provider has been obtained and can be produced upon request.
